# Functional efficacy of the MAO-B inhibitor safinamide in murine substantia nigra *pars compacta* dopaminergic neurons *in vitro*: a comparative study with tranylcypromine

**DOI:** 10.1101/2024.05.28.596142

**Authors:** Beatrice Zarrilli, Cecilia Giacomet, Francesca Cossa, Mauro Federici, Nicola Berretta, Nicola B Mercuri

## Abstract

Safinamide (SAF) is currently used to treat Parkinson’s disease (PD) symptoms based on its theoretical ability to potentiate the dopamine (DA) signal, blocking monoamine oxidase (MAO) B. The present work aims to highlight the functional relevance of SAF as an enhancer of the DA signal, by evaluating its ability to prolong recovery from DA-mediated firing inhibition of DAergic neurons of the substantia nigra pars compacta (SNpc), compared to another MAO antagonist, tranylcypromine (TCP). Using multielectrode array (MEA) and single electrode extracellular recordings of spontaneous spikes from presumed SNpc DAergic cells in vitro, we show that SAF (30 uM) mildly prolongs the DA-mediated firing inhibition, as opposed to the profound effect of TCP (10 uM). In patch-clamp recordings, we found that SAF (30 uM) significantly reduced the number of spikes evoked by depolarizing currents in SNpc DAergic neurons, in a sulpiride (1 uM) independent manner. According to our results, SAF marginally potentiates the DA signal in SNpc DAergic neurons, while exerting an inhibitory effect on the postsynaptic excitability acting on membrane conductances. Thus, we propose that the therapeutic effects of SAF in PD patients partially depends on MAO inhibition, while other MAO-independent sites of action could be more relevant.

## 1. INTRODUCTION

The primary goal in the therapy of Parkinson’s disease (PD) is to compensate for the profound reduction in dopamine (DA) level in the brain that typically occurs in this disease, due to the progressive neurogeneration of DA-releasing neurons in the mesencephalic Sustantia nigra *pars compacta* (SNpc). Historically, the most effective drug used in the treatment of Parkinson’s symptoms is represented by oral L-DOPA, due to its subsequent conversion to dopamine (DA) (Mercuri et al., 2005). The main problem with this therapeutic approach is that its beneficial effects on PD patients are restricted to a so-called “honeymoon period”, after which patients develop motor fluctuations and dyskinesia (Obeso et al., 2000). Several strategies have been implemented to increase and stabilize the DA signal in the brain of PD patients; in particular, inhibitors of monoamine oxidases (MAO) and catechol-O-methyltransferase have been added to the therapeutic arsenal, with the aim to reduce the degradation of DA and prolong its systemic activity.

Among MAO inhibitors used in PD treatment, selegiline was initially introduced, followed by rasagiline, which are both irreversible MAO-B inhibitors also used in the treatment of patients with Alzheimer’s disease (Tzvetkov et al., 2017). Regarding PD, a reversible MAO-B inhibitor has also been introduced, safinamide (Benedetti et al.,1994; Caccia et al., 2006; Binda et al., 2007; Onofrj et al., 2008), although additional sites of action of this drug have been reported, including inhibition of voltage-gated sodium channels (Sciaccaluga et al., 2020).

We have previously investigated the functional effectiveness of MAO inhibitors in potentiating the DA signal in SNpc DAergic neurons *in vitro*. In particular, by exploiting as experimental readout the D_2_ receptor (D_2_R)-mediated firing inhibition that characterizes these neurons (Lacey et al., 1987), we used the degree of prolongation of DA-induced firing inhibition in response to preexposure to various MAO inhibitors, as a powerful and subtle index of their potency (Mercuri et al., 1996). Therefore, to elucidate the functional efficacy of safinamide as MAO inhibitor, we compared its ability to prolog the DA signal, with that of another irreversible MAO inhibitor, tranylcypromine, (Coutts et al., 1987; Ulrich et al., 2017), whose efficacy has been characterized in our laboratories (Mercuri et al., 2000).

## 2. METHODS

### 2.1. Animals

Adult (25 to 40 day-old) male mice C57BL/6J (Charlers River, Cat #632C57BL/6J, RRID:IMSR_JAX:000664) were kept in a temperature controlled environment under a 12 h light/dark cycle and given food and water *ad libitum*. All experiments were carried out in accordance with the European Directive 2010/63/EU, using protocols approved by the Animal Care and Use Committee at the IRCCS Fondazione Santa Lucia (Italy) and by the Italian Ministry of Health (D.lgs 26/2014, authorization n. 876/2020-PR). All efforts were made to minimize the number of animals used and their suffering.

### 2.2. Electrophysiology

#### 2.2.1 Slice preparation

Mice were decapitated under halothane anaesthesia and 250 μm thick horizontal midbrain slices were prepared as previously described (Guatteo et al., 2017). Slices were maintained in an artificial cerebrospinal fluid (ACSF) solution, bubbled with a mixture of O_2_/CO_2_ (95/5%) at 32 °C containing (in mM): 126 NaCl, 2.5 KCl,1.2 MgCl_2_, 1.2 NaH_2_PO_4_, 2.4 CaCl_2_, 10 glucose and 25 NaHCO_3_. The slices were then sequenty transferred to a recording chamber in continuously flowing ACSF (34 °C). All drugs were dissolved in ACSF and bath applied.

#### 2.2.2 Multielectrode array (MEA) recordings

The slice was placed over an 8×8 array of planar electrodes (100 μm interpolar distance and 20 × 20 μm in size) as previously described (Berretta et al., 2010). The voltage signals were filtered (0.1-100 Hz) and digitized (100 kHz) using Mobius software (Alpha MED Sciences, Osaka, Japan, https://www.med64.com/products/med64-mobius-software/). When needed, spike-sorting was achieved according to spike waveform with a normal mixtures algorithm on independent clusters obtained from principal component data (Spike2 ver. 6.0 software, RRID:SCR_000903, Cambridge Electronic Design Ltd, Cambridge, UK, http://www.ced.co.uk/pru.shtml?spk7wglu.htm) (Berretta et al., 2010; Aversa et al., 2018).

#### 2.2.3 Single-electrode single-unit recordings

Spontaneous extracellular single-units were recorded from midbrain slices placed in a submerged chamber. The glass microelectrodes had a tip resistance of 5–10 MΩ when filled with ACSF. The recordings were made with an Axoclamp 900A amplifier using Clampex 10 software and digitized with a Digidata 1440 (Molecular Devices, LLC., 3860 N First Street San Jose,CA 95134 USA).

#### 2.2.4 Patch-clamp recordings

Whole-cell patch-clamp recordings were performed with borosilicate glass pipettes from SNpc neurons, visualized using infrared microscopy (Nikon, Stroombaan 14, 1181 VX Amstelveen, The Netherlands), using a Multiclamp 700B amplifier (Molecular Devices, LLC., 3860 N First Street San Jose, CA 95134 USA). The standard internal solution contained (in mM): 135**□**K-gluconate, 10 KCl, 10 HEPES, 2 MgCl_2_, 0.1 EGTA, 0.05 CaCl_2_, 4 Mg_2_-ATP, 0.3 Na_3_-GTP, 10 Phosphocreatine-Na_2_, adjusted to pH 7.3 with KOH.

Recorded neurons were classified as DA neurons based on electrophysiological criteria, i.e. the presence of Ih in response to hyperpolarizing voltage steps in voltage-clamp mode (Krashia et al., 2017). The protocol used to measure evoked action potential firing in current-clamp mode consisted of 2-s depolarizing current steps (from +50 to +250**□**pA, 50**□**pA increasing steps) from an imposed resting membrane potential of -60 mV, by constant current injection.

### 2.3 Drugs

Safinamide, tranylcypromine and L-sulpiride (all from Merck KGaA, Darmstadt, Germany) were dissolved in ACSF and bath applied at their final concentration by a three-ways tap.

### 2.4 Statistics

Statistical analyses were performed with OriginPro software ver. 2024 (RRID:SCR_014212, Northampton, MA 01060 USA, http://www.originlab.com/index.aspx?go=PRODUCTS/Origin). For both MEA and single-electrode single-unit recordings, the firing rate of each neuron was normalized to its control level and avareged across neurons. According to the Kolgomorov-Smirnov normality test the data were not normally distributed, thus, for comparisons we performed the Mann-Whitney test and no correction for multiple comparisons was used. For patch-clamp recordings, we first assessed data normality by using the Kolgomorov-Smirnov normality test. Then, two-way repeated measures (RM) analysis of variance (ANOVA) with Tukey’s post-hoc comparison was applied to normally distributed data, while Friedman ANOVA followed by paired sample Wilcoxon signed ranks test was used when data were not normally distributed.

## 3. RESULTS

### 3.1. Comparison between SAF and TCP on DA-mediated firing inhibition of SNpc DAergic neurons

#### 3.1.1 Multi-electrode array recordings

We recorded the spontaneous firing of presumed DAergic neurons located in the SNpc, identified according to the presence of inhibition of the spontaneous firing in response to a brief perfusion with DA (30 μM, 3 min), due to their typical D_2_R-inhibitory autoreceptors (Lacey et al., 1987). The time for recovery of this inhibitory response at DA washout was then compared to that obtained after superfusion in 30 μM safinamide (SAF), a concentration previously shown to exert maximal effects as MAO inhibitor in slice preparations (Sciaccaluga et al., 2010).

SAF perfusion for 30 min did not modify on its own the basal firing rate (p > 0.1 Mann-Whitney U test, n=29 cells from 6 slices; data not shown), however, DA-mediated firing inhibition was more profound and long-lasting than under control conditions, before recovering to baseline after 15 min from DA washout (p < 0.05 Mann-Whitney U test, n=29 cells from 6 slices; Fig. 1A). More prolonged exposures in 30 μM SAF at 60 or 120 min still resulted in a higher and longer DA-mediated firing inhibition (p < 0.001 Mann-Whitney U test, n=58 cells from 15 slices after 60 min and p < 0.005 Mann-Whitney U test, n=37 cells from 8 slices after 120 min; Fig. 1A), although not significantly different from that observed after 30 min SAF (p > 0.1 Mann-Whitney U test), indicating that a maximal effect was already achieved with this exposure time.

**Fig. 1.**
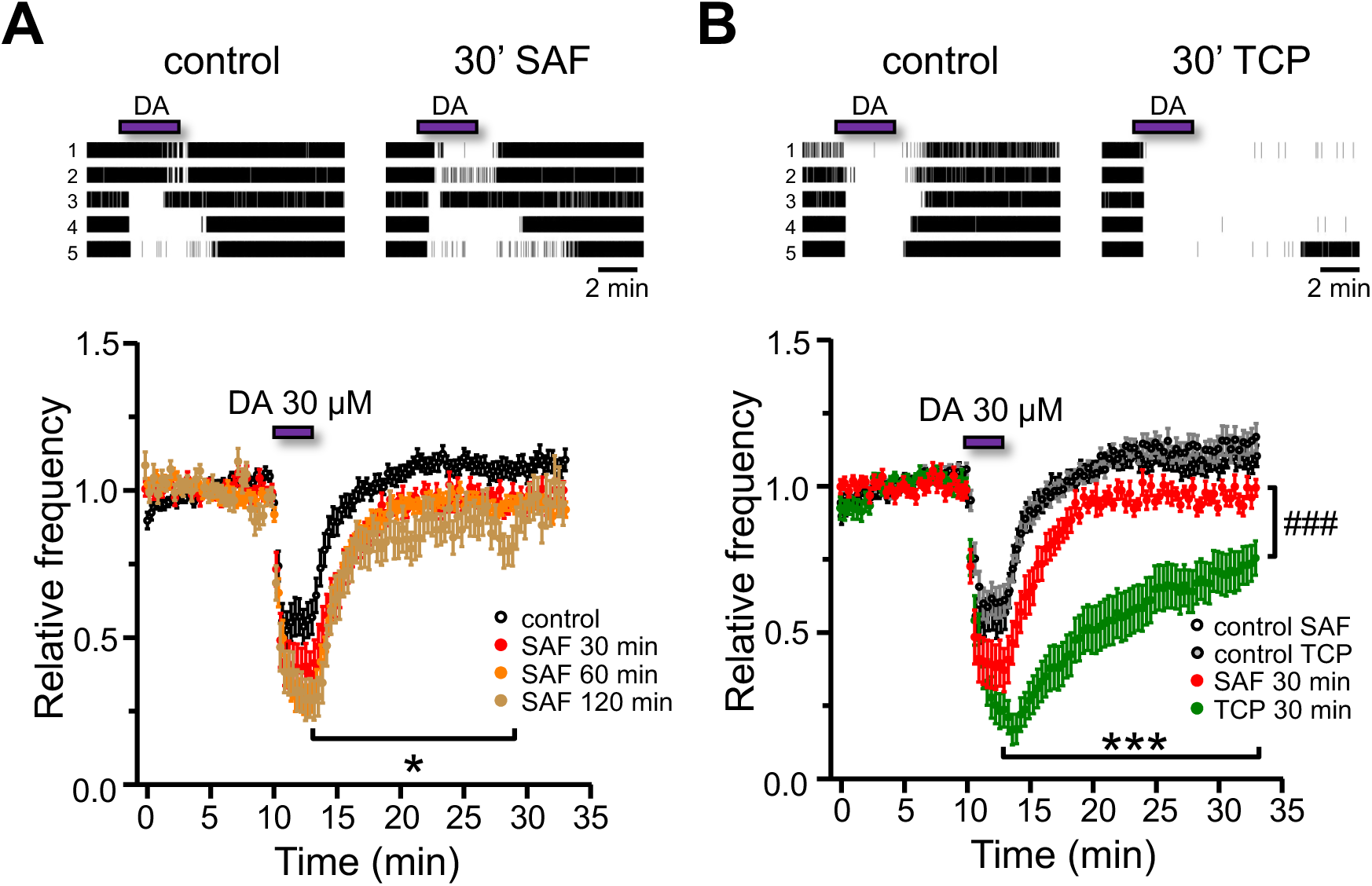
Effect of safinamide and tranylcypromine on the firing of SNpc DAergic neurons recorded with MEA. Running average of the mean (± ES) firing frequency (bin size 20s) of SNpc neurons recorded with MEA, normalized to the 10 min preceding each 3 min exposure to DA (30 μM). (A) The recovery rate from DAmediated firing inhibition was significantly reduced after 30, 60 and 120 min in SAF (30 μM), reaching a maximal effect after 30 min (*, p < 0.05 Mann-Whitney U test). On top, raster plots of the firing of 5 selected neurons in control and in SAF (30 μM). (B) After 30 min perfusion in TCP (10 μM) the recovery rate from DA-mediated firing inhibition was significantly reduced compared to control (***, p < 0.001 Mann-Whitney U test). The effect on the firing rate is also shown in 30 μM SAF with its respective control (same data of panel A), showing the significantly larger effect of TCP compared to SAF (###, p < 0.001 Mann-Whitney U test). On top, raster plots of the firing of 5 selected neurons in control and in TCP (10 μM).

We then compared the effect of 30 min SAF (30 μM) on the degree of DA-mediated firing inhibition, with that of another MAO inhibitor, tranylcypromine (TCP), at a concentration of 10 μM, previously shown to exert maximal effects *in vitro* (Mercuri et al., 2000). As expected, 30 min exposure in 10 μM TCP significantly increased firing inhibition by 30 μM DA (p < 0.001 Mann-Whitney U test, n=38 cells from 5 slices; Fig. 1B). Furthermore, the degree and duration of DA-mediated firing inhibition was significantly higher than that observed after 30 μM SAF for 30 min (p < 0.001 Mann-Whitney U test), indicating a lower functional efficacy of SAF in increasing the DA signal compared to TCP.

#### 3.1.2 Single electrode single-unit recordings

The effect of safinamide was further evaluated by conventional extracellular recordings of spontaneous action potentials with sharp microelectrodes from individual SNpc DAergic neurons, identified according to occurrance of firing inhibition in response to DA (30 μM, 1.5 min).

Pretreatment in SAF (30 μM) for 30 min did not modify the peak of DA-mediated firing inhibition, compared to control, however a significant delay in the recovery was observed, before the firing rate returned to control levels after 20 min from DA washout (p < 0.05 Mann-Whitney U test, n=10 cells; Fig. 2A).

**Fig. 2.**
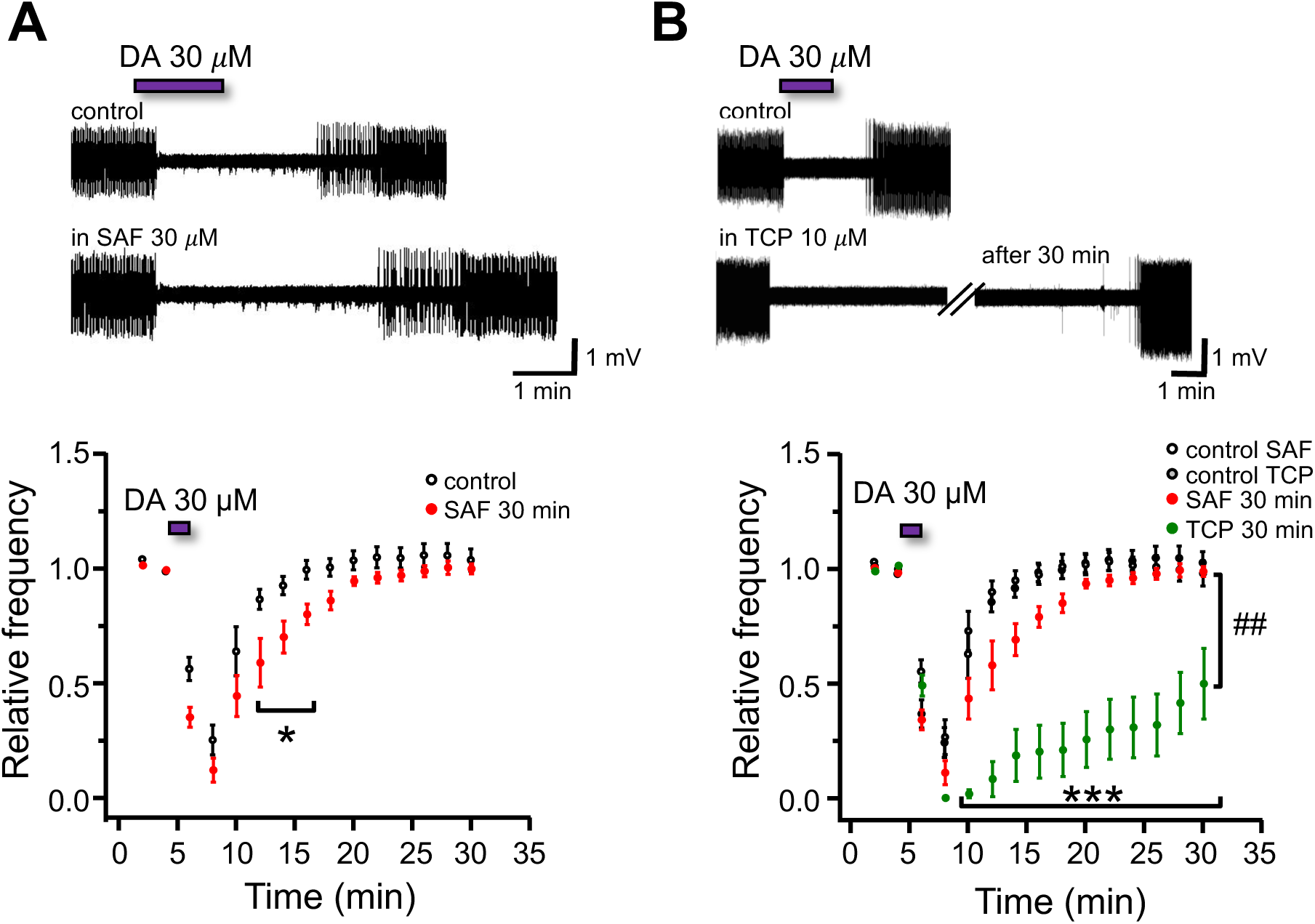
Effect of safinamide and tranylcypromine on the firing of SNpc DAergic neurons recorded with extracellular sharp electrodes. Running average of the mean (± ES) firing frequency (bin size 60s) of SNpc neurons recorded with extracellular sharp electrodes, normalized to the 2 min preceding each 1.5 min exposure to DA (30 μM). (A) After 30 min in SAF (30 μM), the recovery rate from DA-mediated firing inhibition was significantly reduced compared to control (*, p < 0.05 Mann-Whitney U test). On top, sample traces of extracellular action potentials from a single neuron, in control and 30 min after exposure of SAF 30 μM. (B) After 30 min perfusion in TCP (10 μM) the recovery rate from DA-mediated firing inhibition was significantly reduced compared to control (***, p < 0.001 Mann-Whitney U test). The effect on the firing rate is also shown in 30 μM SAF with its respective control (same data of panel A), showing the significantly larger effect of TCP compared to SAF (##, p < 0.01 Mann-Whitney U test). On top, sample traces of extracellular action potentials from a single neuron, in control and 30 min after exposure of TCP 10 μM.

When we measured the effect of 10 μM TCP, under similar experimental conditions, DA-induced firing inhibition was more profound and lasting compared to control (p < 0.001 Mann-Whitney U test, n=8 cells; Fig. 2B), with an overall effect of TCP that was significantly more pronounced than that of 30 μM SAF (p < 0.01 Mann-Whitney U test).

#### 3.1.3 Effect of Safinamide on DAergic neurons’ excitability

Previous studies indicate that SAF not only acts as MAO inhibitor, but it also affects neuronal excitability through inhibition of voltage-gated conductances (Sciaccaluga et al., 2020). To verify the presence of a similar mechanism of action on SNpc DAergic neurons, we tested if SAF affected the current-driven firing discharge of DAergic cells recorded in the whole-cell patch-clamp.

Neuronal firing was induced with 2-s depolarizing current steps (from +50 to +250 pA, 50**□**pA increasing steps) from an imposed resting membrane potential of -60 mV. This protocol was used in control and after 10 to 12 min of SAF 30 μM perfusion. We observed that SAF reduced the number of action potentials evoked with higher depolarizing steps (200 to 250 pA; p < 0.01 paired sample Wilcoxon signed ranks test; n = 10; Fig 3A).

**Fig. 3.**
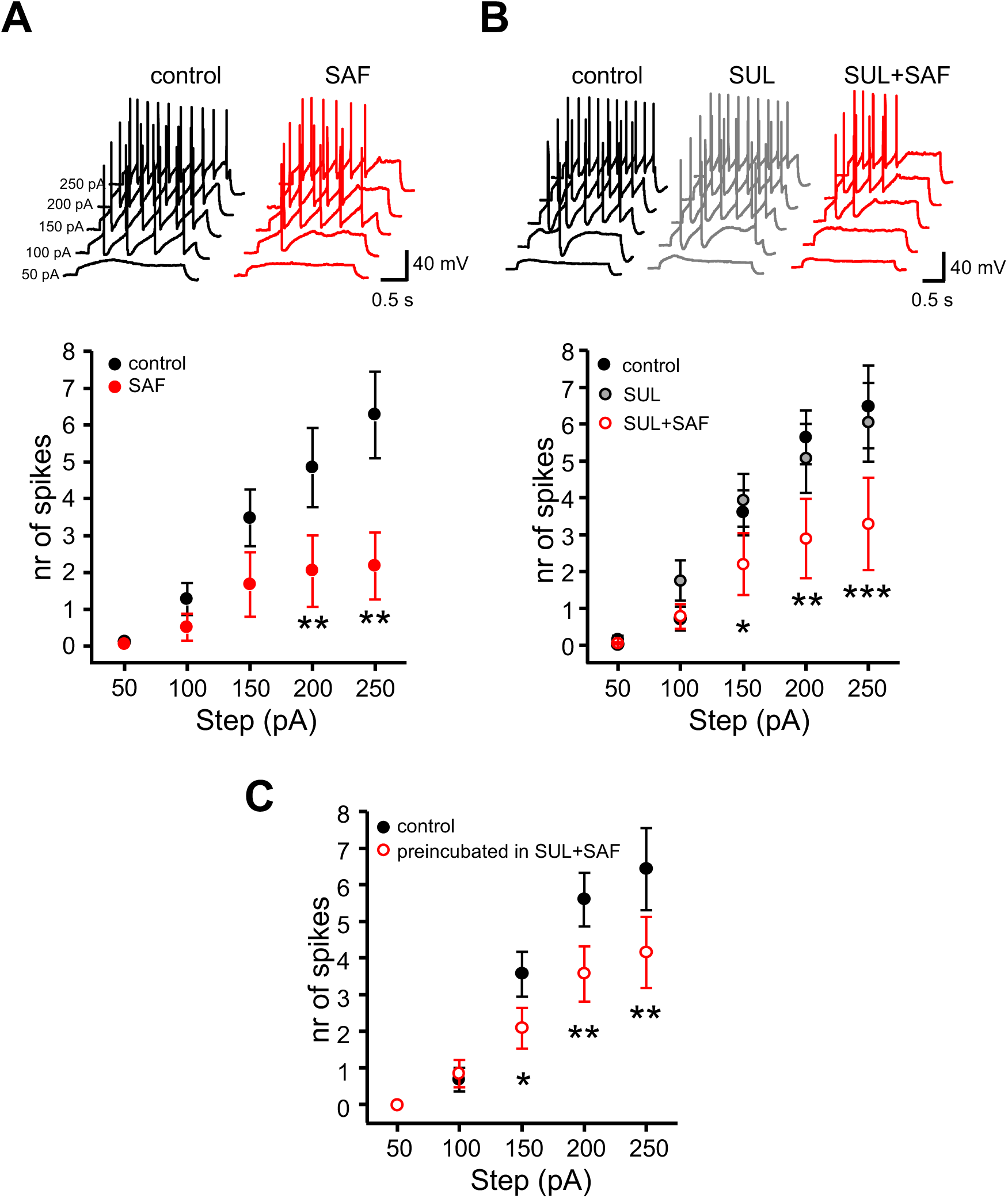
Effect of safinamide on the evoked firing of SNpc DAergic neurons recorded with patch-clamp technique. Data are represented as mean (± SEM) of the number of action potentials for each depolarizing step (50 to 250 pA, 50 pA increment). (A) The number of evoked spikes was significantly reduced after the treatment in SAF (30 μM) (Friedman ANOVA: control p < 0.0001, SAF p = 0.077; **, p < 0.01 paired sample Wilcoxon signed ranks test at 200 and 250 pA). On top, representative traces of the evoked spikes in the same cell before and after treatment with SAF (30 μM). (B) Treatment in SAF (30 μM) significantly reduced the number of evoked spikes also in the presence of SUL (1 μM) (*, p < 0.05 at 150 pA; **, p < 0.01 at 200 pA; ***, p < 0.001 at 250 pA, two-way ANOVA with Tukey’s post hoc comparison between SUL only and SUL+SAF treatments; p < 0.001 at 200 pA and 250 pA, two-way ANOVA with Tukey’s post hoc comparison between control and SUL+SAF treatment). On top, representative traces of the evoked spikes in the same cell before and after treatment with SUL (1 μM) alone or with added and SAF (30 μM). (C) SNpc DA neurons in SUL (1 μM) and SAF (30 μM) pretreated slices show a significant reduction in the number of evoked action potentials in comparison to neurons in untreated slices (same data of panel A) (*, p < 0.05 at 150 pA; **, p < 0.01 at 200 pA and at 250 pA, two-way ANOVA with Tukey’s post hoc comparison between untreated and SUL+SAF treated cells).

To assess if this decreased excitability by SAF was due to a DA-dependent mechanism, through stimulation of postsynaptic D_2_R, we evaluated whether treatment with the D_2_R antagonist sulpiride 1 μM (SUL) could prevent the effect of SAF on membrane excitability. Thus, we pretreated the same neurons for 10 min in 1 μM SUL and repeated SAF (30 μM) perfusion in the presence of SUL. Under these conditions, postsynaptic excitability was still reduced at higher depolarizing steps (150 pA, p < 0.05; 200 pA, p < 0.01; 250 pA, p < 0.001 two-way ANOVA with Tukey’s post hoc comparison; n = 10; Fig. 3B). This result was confirmed by comparing the cellular excitability in separate DA neurons recorded in control slices or in slices preincubated in SAF 30 μM and SUL 1 μM. This protocol was used to rule out that the observed reduction in excitability might be due to prolonged dialysis of the patched neurons that have undergone successive SUL and SAF treatments. Also in these experimental conditions, the number of evoked action potentials was higher in neurons recorded from slices maintained in ACSF than in neurons of slices pretreated in SAF and SUL (150 pA, p < 0.05; 200 to 250 pA, p < 0.01 two-way ANOVA with Tukey’s post hoc comparison; n = 10 control vs 12 treated cells; Fig. 3C).

## 4. DISCUSSION

By performing electrophysiological recordings of SNpc neurons in midbrain slices, we have shown that SAF, a selective and reversible monoamine oxidase-B inhibitor (Benedetti et al.,1994; Caccia et al., 2006; Binda et al., 2007; Onofrj et al., 2008) mildly prolongs the DA signal, measured by the extent of firing inhibition of SNpc DAergic neurons in response to DA. Despite the saturating concentration of SAF used (30 μM; see Sciaccaluga et al., 2020), this effect was significantly less pronounced than that of TCP, another well-known MAO blocker, which instead produces a prolonged inhibition of neuronal firing in response to DA, in analogy with previous observations (Mercuri et al., 2000).

The limited functional effect of SAF compared to TCP could be due to the need to block both MAO B and A, because DA is a substrate for both isoforms (Geha et al., 2001). Indeed, while SAF is selective for MAO-B, TCP has been reported to block MAO A and B (Ulrich S et al., 2017). In line with this, concerning other amines, a combination of MAO-A and MAO-B inhibitors (clorgyline plus L-deprenyl) is more effective in increasing brain serotonin and noradrenaline (Green and Youdim, 1975).

Our results also show that SAF significantly reduces the postsynaptic excitability of SNpc DAergic neurons by reducing the number of action potentials evoked by a depolarizing current. This effect cannot be ascribed to increased extracellular DA acting on inhibitory D_2_ autoreceptors, as the reduced excitability was still observed in the presence of the D_2_R antagonist sulpiride, thereby pointing to an effect on voltage-dependent membrane conductances. In fact, previous evidence in other neurons has demonstrated a direct effect of SAF on sodium and calcium voltage-gated channels (Salvati et al., 1999; Sciaccaluga et al., 2020). This latter mechanism of action could regulate excitatory neurotransmitter release in the basal ganglia (Gardoni F et al., 2018 ; Muller and Foley, 2017), thus reducing motor fluctuations in response to L-DOPA (Kandadai et al., 2014; Schapira et al., 2017; Bhidayasiri et al., 2023). In PD patients, administration of SAF together with L-DOPA increases ‘on’ period duration and reduces adverse symptoms in ‘off’ period which might principally correlate with a regulation of synaptic plasticity in the basal ganglia (Sciaccaluga et al., 2020) via reduction of the abnormal release of glutamate in the DA-denervated state. A regulation of neurotransmitter release by SAF is also supported by the present electrophysiological data, where no potentiation of the DAergic signal has been detected.

Thus, we suggest that other actions unrelated to MAO B and DA interaction should preferentially account for the therapeutic effects of safinamide in Parkinson’s disease patients.

## DATA AVAILABILITY

Full datasets can be found at Zenodo repository: 10.5281/zenodo.11262613. A detailed description of the conducted protocol is available at protocols.io repository: dx.doi.org/10.17504/protocols.io.e6nvw1jwzlmk/v1

## ACKNOWLEDGMENTS

This research was funded in whole or in part by Aligning Science Across Parkinson’s [grant ASAP-020505] through the Michael J. Fox Foundation for Parkinson’s Research (MJFF). For open access, the author has applied a CC BY public copyright license to all Author Accepted Manuscripts arising from this submission.

